# Targeting repair pathways with small molecules increases precise genome editing in pluripotent stem cells

**DOI:** 10.1101/136374

**Authors:** Stephan Riesenberg, Tomislav Maricic

## Introductory paragraph

A now frequently used method to edit mammalian genomes uses the nucleases CRISPR/Cas9 and CRISPR/Cpf1 or the nickase CRISPR/Cas9n to introduce double-strand breaks (DSBs) which are then repaired by homology-directed repair (HDR) using synthetic or cloned DNA donor molecules carrying desired mutations. However, another pathway, the non-homologous end joining (NHEJ) pathway competes with HDR for repairing DNA breaks in cells. To increase the frequency of precise genome editing in human induced pluripotent stem cells (hiPSCs) and human embryonic stem cells (hESCs) we have tested the capacity of a number of small molecules to enhance HDR or inhibit NHEJ. We identify molecules that increase the frequency of precise genome editing including some that have additive effects when applied together. Using a mixture of such molecules, the ‘CRISPY’ mix, we achieve 2.8- to 6.7-fold increase in precise genome editing with Cas9n, resulting in the introduction of the intended nucleotide substitutions in almost 50% of chromosomes, to our knowledge the highest editing efficiency in hiPSCs described to date. Furthermore, the CRISPY mix improves precise genome editing with Cpf1 2.9- to 4.0-fold, allowing almost 20% of chromosomes to be edited.

## Main text

The bacterial nuclease CRISPR/Cas9 is now frequently used to accurately cut chromosomal DNA sequences in eukaryotic cells. The resulting DNA breaks are repaired by two competing pathways: NHEJ and HDR (Fig. 1). In NHEJ, the first proteins to bind the cut DNA ends are Ku70/Ku80, followed by DNA protein kinase catalytic subunit (DNA-PKcs)^1^. The kinase phosphorylates itself and other downstream effectors at the repair site which results in joining of the DNA ends by DNA ligase IV^2^. If this canonical NHEJ is repressed, the alternative NHEJ pathway becomes active^3^ which, amongst other proteins, requires Werner syndrome ATP-dependent helicase. HDR is initiated when the MRN complex binds to the DSB^1^. In this case, DNA endonuclease RBBP8 (CtIP) removes nucleotides at the 5' ends. Further resection produces long 3' single-stranded DNA (ssDNA) overhangs on both sides of the DNA break^2^. These are coated and stabilized by the replication protein A (RPA) complex, followed by generation of a Rad51 nucleoprotein filament^1^. Rad52 facilitates replacement of RPA bound to ssDNA with Rad51 and promotes annealing to a homologous donor DNA^4^. Subsequent DNA synthesis results in precisely repaired DNA. In HDR, the protein kinase ataxia-telangiectasia mutated (ATM) plays a major role in that it phosphorylates at least 12 proteins involved in the pathway^1^.

**Fig. 1:**
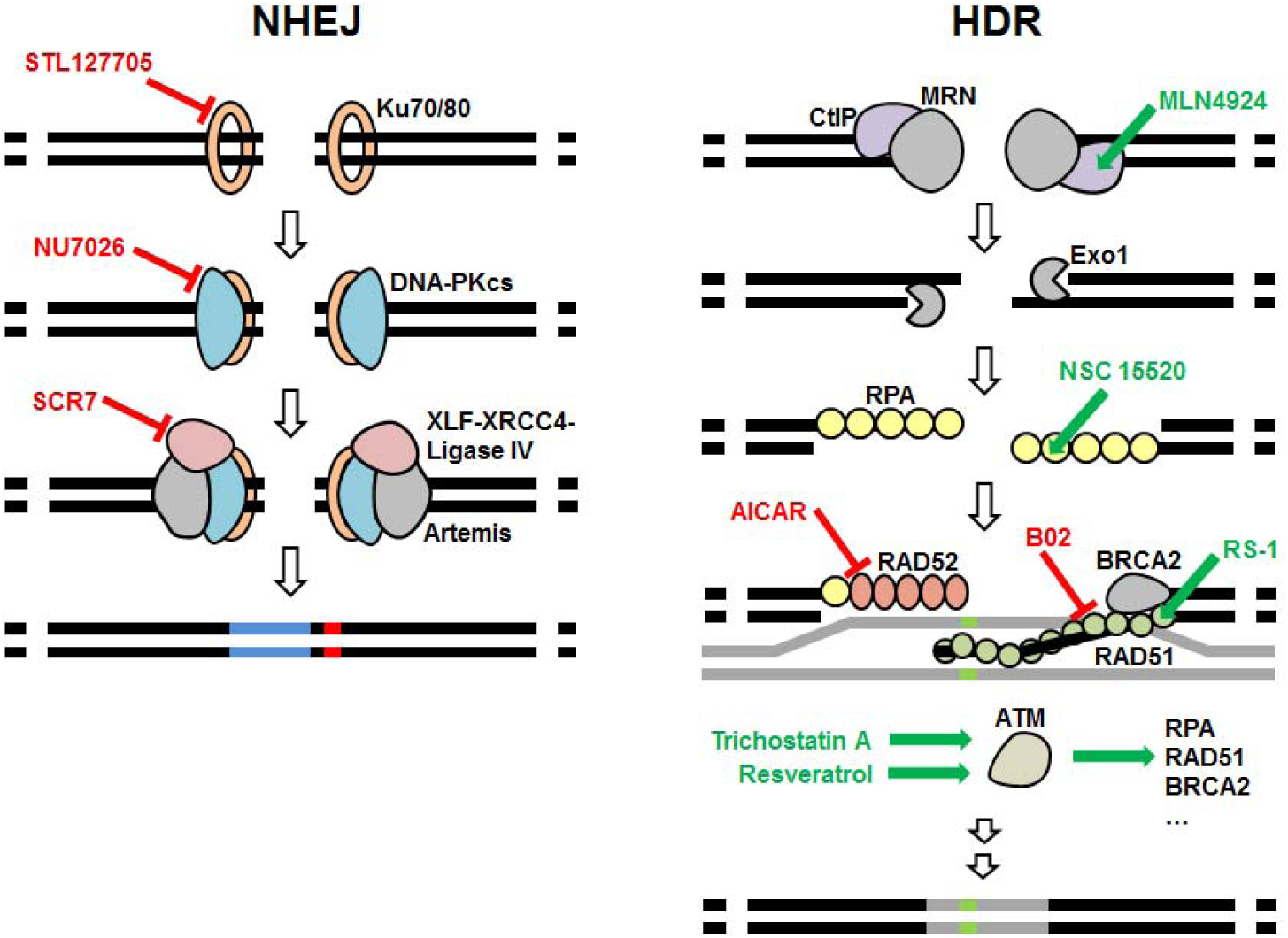
Small molecules described or anticipated to target key proteins of NHEJ and HDR. Proteins, are labelled with black text and inhibitors and enhancing small molecules are marked red and green, respectively. STL127705, NU7026 or SCR7 have been described to inhibit Ku70/80, DNA-PK or ligase IV, respectively. MLN4924, RS-1, Trichostatin A or Resveratrol have been described to enhance CtIP, RAD51 or ATM, respectively. NSC 15520 has been described to block the association of RPA to p53 and RAD9. AICAR is an inhibitor of RAD52 and B02 is an ihibitor of RAD51.

Since NHEJ of Cas9-induced DSBs is error prone and frequently introduces short insertions and deletions (indels) at the cut site, it is useful for knocking out a targeted gene. In contrast, HDR allows precise repair of a DSB by using a homologous donor DNA. If the donor DNA provided in the experiment carries mutations, these will be introduced into the genome (precise genome editing). Repair with homologous ssDNA or dsDNA has been suggested to engage different pathways^5^. We will refer to Targeted Nucleotide Substitutions using ssDNA donors as ‘TNS’ and targeted insertion of cassettes using dsDNA donors as knock-ins, respectively. In order to introduce a DSB Cas9 requires the nucleotide sequence NGG (a “PAM” site) in the target DNA. Targeting of Cas9 is further determined by a guide RNA (gRNA) complementary to 20 nucleotides adjacent to the PAM site. However, the Cas9 may also cut the genome at sites that carry sequence similarity to the gRNA^6^. One strategy to reduce such off-target cuts is to use a mutated Cas9 that introduces single-stranded nicks instead of DSBs (Cas9n)^7^. Using two gRNAs to introduce two nicks on opposite DNA strands in close proximity to each other will result in a staggered DSB at the desired location while reducing the risk of off-target DSBs because two nicks close enough to cause a DSB are unlikely to occur elsewhere in the genome. Another strategy is to use Cpf1^8^, a nuclease that introduces staggered cuts near T-rich PAM sites and causes less off-target DSBs than Cas9^9, 10^.

Efficiencies of TNS in human stem cells range from 15% down to as low as 0.5%^11, 12^ making the isolation of edited homozygous clones challenging. Several studies have tried to increase precise genome editing efficiency by promoting HDR or decreasing NHEJ. Synchronization of cells to the S or G_2_/M phase when homologous recombination occurs increases TNS efficiency in HEK cells (from 26% to 38%), human primary neonatal fibroblast (undetectable to 0.6%) and hESCs (undetectable to 1.6%)^13^ and knock-in efficiency in hESCs (from 7 to 41%)^14^. Improved knock-in efficiency was also achieved in HEK cells by suppressing repair proteins like Ku70/80 and DNA ligase IV with siRNA (from 5 to 25%) or by coexpression of adenovirus type 5 proteins 4E1B55K and E4orf6 which mediate degradation of DNA ligase IV among other targets (from 5 to 36%)^15^.

Several small molecules have been used to increase precise genome editing (Supplementary Table 1). DNA ligase IV inhibitor SCR7 has been claimed to block NHEJ and increase the efficiency of TNS (from 5 to 22.7%) in mouse embryos^16^. Others described an increase of knock-in efficiency in HEK cells^15, 17^, while others found no significant effect in rabbit embryos^18^ and human stem cells^14, 19^. Recently, Greco et al. reanalysed the structure and inhibitory properties of SCR7^20^ and concluded that SCR7 and its derivates are neither selective nor potent inhibitors of human DNA ligase IV. Inhibition of DNA-PK in the NHEJ-pathway by the small molecules NU7441 and NU7026 has been shown to reduce the frequency of NHEJ and to increase efficiency of knock-ins in HEK cells (from 1.9 to 3.8%; from 3 to 7.6%)^21, 22^ and hiPSCs (from 13 to 16%)^19^ and of TNS in mouse embryonic fibroblasts (from 3 to 10%)^21^. The RAD51 stimulatory compound RS-1 increased knock-in efficiency in rabbit embryos (from 4.4 to 26.1%)^18^, HEK cells (from 3.5 to 21%)^17^ and U2OS cells (from 1.9 to 2.4%)^17^, but not in hiPSCs^19^. No effect of RS-1 on TNS efficiency was found in porcine fetal fibroblasts^23^. Furthermore, using a library screen of around 4,000 small molecules, Yu et al. found the β3-adrenergic receptor agonist L755507 to increase TNS efficieny in hiPSCs (from 0.35 to 3.13%) and knock-in efficieny in mouse ESCs (from 17.7 to 33.3%), although it is not know how this molecule interacts with repair pathways^12^. Others did not find any significant stimulation of HDR by L755507 in HEK cells^17^ or hiPSCs^19^. Pinder et al. used SCR7, RS-1 and L755507 singly and together and found no additive effect when using either SCR7 or L755507 together with RS-1 compared to RS-1 alone. In summary, inhibitors of DNA-PK increase precise genome editing efficieny in different cell lines while the effects of SCR7, L755507 and RS-1 are not consistent between cell lines.

Here we test the above as well as other small molecules with respect to their efficiency to induce TNS in hiPSCs. We indentified additional molecules interacting with repair proteins listed in the REPAIRtoire database^24^ by literature and database (ChEMBL^25^) search. The additional molecules we test, which have been described to block NHEJ or alternative NHEJ or activate or increase the abundance of proteins involed in HDR or damage-dependent signaling (Fig. 1 and Supplementary Table 2), are: NU7026, Trichostatin A, MLN4924, NSC 19630, NSC 15520, AICAR, Resveratrol, STL127685 and B02.

We tested these molecules in iPSC lines that we generated that carry doxycycline-inducible Cas9 (iCRISPR-Cas9) and Cas9 nickase with the D10A mutation (iCRISPR-Cas9n) integrated in their genomes^11^. After the delivery of gRNA and ssDNA donor molecules, cells were treated with small molecules for 24 hours, expanded, their DNA was collected, targeted loci sequenced and editing efficiency quantified (Supplementary Fig. 1, Supplementary Fig. 2).

We tested the effect of different concentrations of each molecule on TNS in the three genes *CALD1, KATNA1* and *SLITRK1* in iCRISPR Cas9n cells. For further experiments, we used the concentration that gave the highest frequency of TNS or if two or more concentrations gave a similarly high frequency we chose the lowest concentration (Supplementary Fig. 3). We found that NU7026 increased TNS 1.5-fold for *CALD1*, 2.6-fold for *KATNA1* and 2.5-fold for *SLITRK1* in Cas9n cells (Fig. 2A), and 1.5-fold for *CALD1*, 1.6-fold for *KATNA1* and 1.2-fold for *SLITRK1* in Cas9 cells (Fig. 2B). Trichostatin A and MLN4924 increased TNS in Cas9n cells 1.5-fold for *CALD1*, 2.2-fold for *KATNA1* and 1.8-fold for *SLITRK1* while no increase was seen in Cas9 cells. MLN4924 increased TNS 1.2-fold for *CALD1*, 1.1- fold for *KATNA1* and 1.3-fold for *SLITRK1* in Cas9n cells while it slightly reduced TNS in Cas9 cells. NSC 15520 increased TNS of *CALD1* 1.4-fold and 1.3-fold in Cas9n and Cas9 cells, respectively, but had no effect on TNS of *KATNA1* and *SLITRK1*. NSC 19630, AICAR, RS-1, Resveratrol, SCR7, L755507 and STL127685 showed no clear effect on TNS frequency in the three genes in Cas9n cells and had no effect or decreased TNS in Cas9 cells. B02 reduced TNS in all three genes in both cell lines (Fig. 2).

**Fig. 2:**
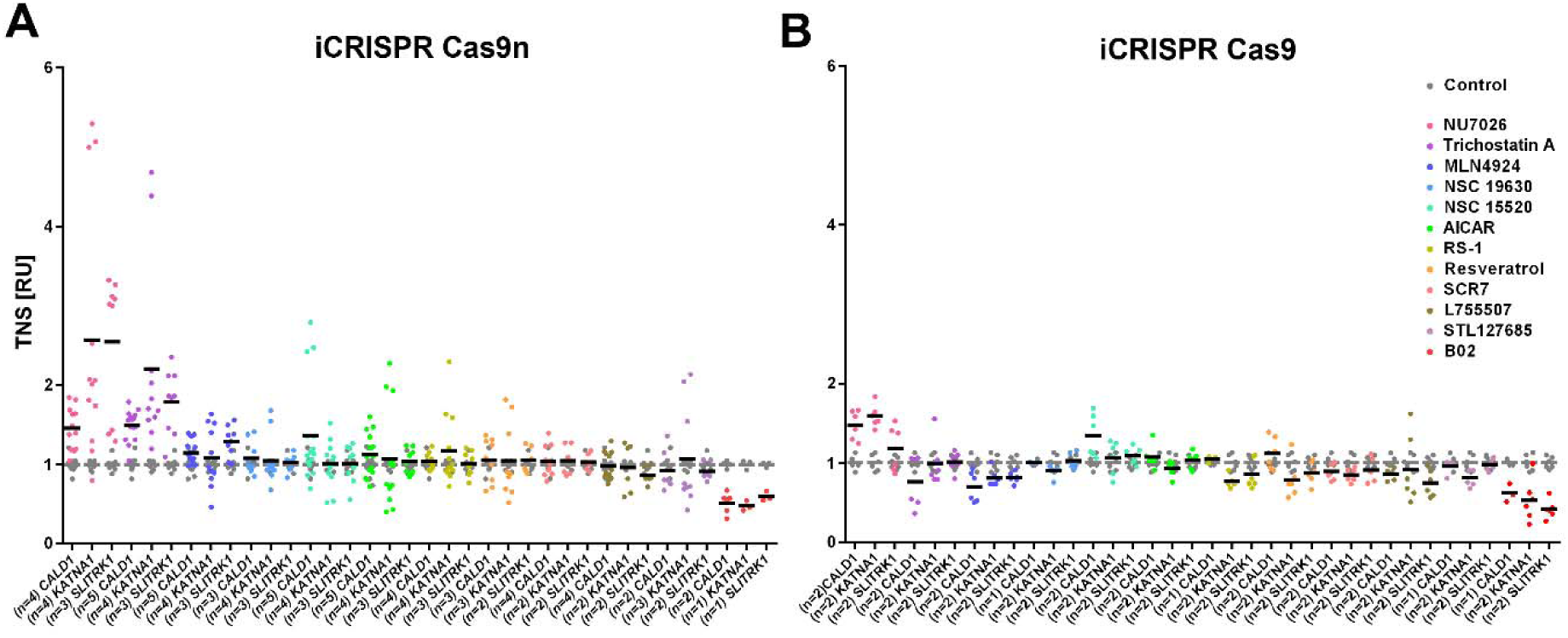
Effects of small molecules on Targeted Nucleotide Substitution (TNS) efficiency in *CALD1*, *KATNA1* and *SLITRK1* with Cas9n and Cas9. TNS efficiency is given in relative units (RU) with the mean of controls set to 1 to account for varying efficiency in different loci. Shown are technical replicates of n independent experiments. Data from Supplementary Fig. 1 and Fig. 2 are included. Grey and black bars represent the mean of the control and the respective small molecule, respectively. Concentrations used were 20μM NU7026, 0.01μM Trichostatin A, 0.05μM MLN4924, 1μM NSC 19630, 5μM NSC 15520, 20μM AICAR, 1μM RS-1, 1μM Resveratrol, 1μM SCR7, 5μM L755507, 5μM STL127685 and 20μM B02.

To test if combinations of these compounds enhance TNS we combined compounds that individually increased TNS for at least one gene in Cas9n cells and never decreased TNS. Those are NU7026, Trichostatin A, MLN4924, NSC 19630, NSC 15520, AICAR and RS-1. The results are shown in Fig. 3A and B. Treatment with NU7026 or Trichostatin A resulted in 2.3- or 1.8-fold higher TNS in Cas9n cells (Tukey's pair-wise post-hoc comparisons: p<0.001) (Fig. 3A) and combinations of NU7026 and Trichostatin A resulted in 1.3 to 1.6 times higher TNS than with either compound alone (p<0.001). Addition of MLN4924 to the mix of NU7026 and Trichostatin A lead to an additional 1.3-fold increase in TNS (p<0.01). Further addition of NSC 15520 slightly increased the mean TNS in Cas9n cells, without reaching statistical significance. Addition of NSC 19630, AICAR and RS-1 had no measurable effect on TNS. We conclude that the mix of small molecules that increases the frequency of TNS with Cas9n the most (although we admittedly could not test all combinatorial possibilities) is a combination of NU7026 (20μM), Trichostatin A (0.01μM), MLN4924 (0.5μM) and NSC 15520 (5μM). This ‘CRISPY’ nickase mix results in an increase of TNS of 2.8-fold (from 11 to 31%) for *CALD1*, 3.6-fold (from 12.8 to 45.8%) for *KATNA1* and 6.7-fold (from 4.7 to 31.6%) for *SLITRK1* in the iCRISPR 409- B2 iPSC line.

**Fig. 3:**
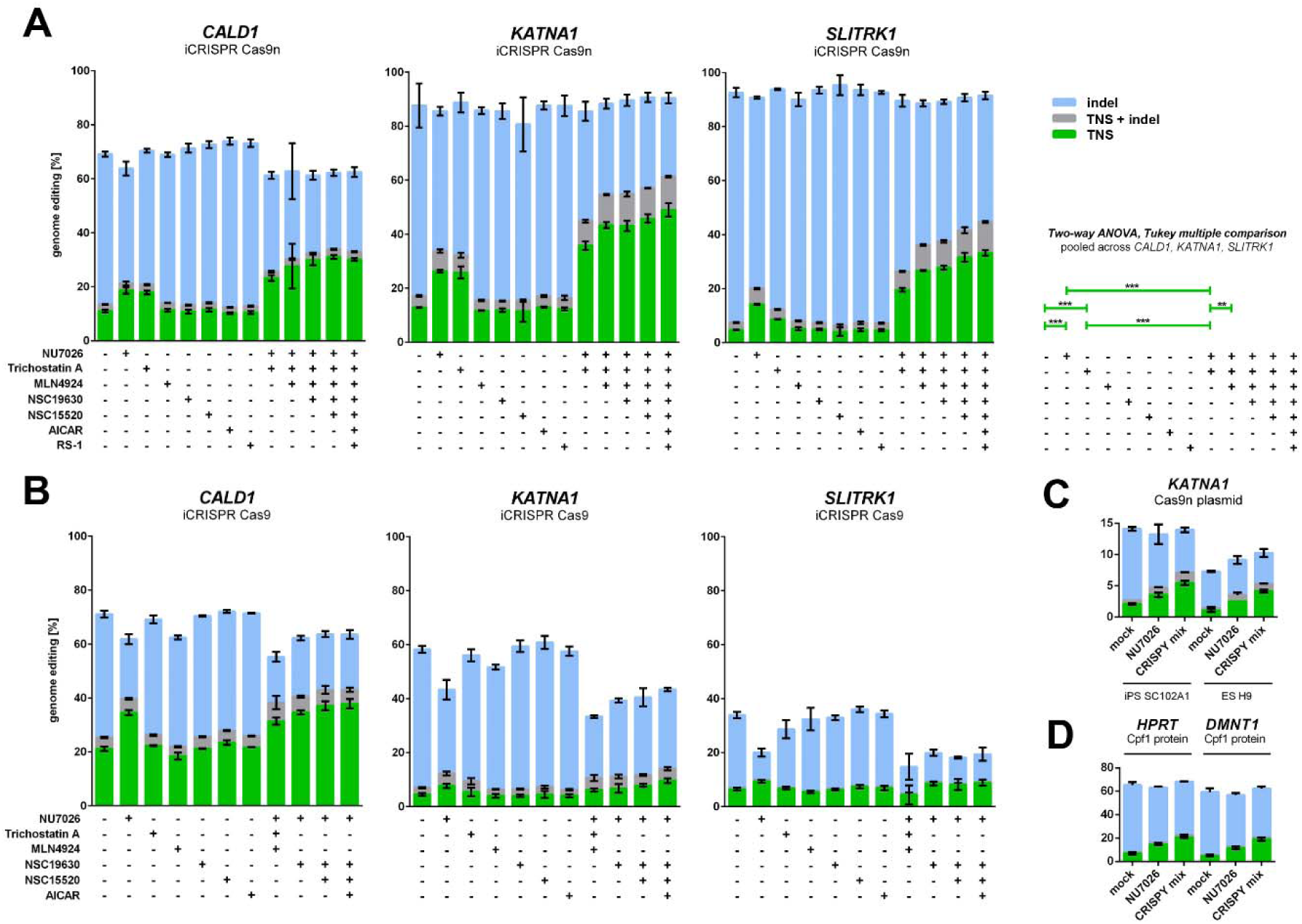
Impact of small molecule combinations on Targeted Nucleotide Substitution (TNS) efficiency in *CALD1*, *KATNA1* and *SLITRK1* with Cas9n and Cas9, and in *HPRT* and *DMNT1* with Cpf1. Small molecules have an additive effect on TNS efficiency with Cas9n (A) but not with Cas9 (B) in the 409-B2 iCRISPR iPSC lines. TNS efficiency of *KATNA1* was also increased in SC102A1 iPSCs and H9 ESCs using the CRISPY mix with plasmid-delivered Cas9n (C). TNS of *HPRT* and *DMNT1* in 409-B2 iPSCs with recombinant Cpf1 was increased using the CRISPY mix as well (D). Shown are TNS, TNS + indels, and indels with green, grey or blue bars, respectively. Error bars show the standard deviation of three technical replicates for A,B and D and two technical replicates for C. Concentrations used were 20μM of NU7026, 0.01μM of Trichostatin A, 0.05μM MLN4924, 1μM NSC 19630, 5μM NSC 15520, 20μM AICAR and 1μM RS-1. CRISPY mix indicates a small molecule mix of NU7026, Trichostatin A, MLN4924 and NSC 15520. Statistical significances of TNS efficiency changes was determined using a two-way-ANOVA and Tukey multiple comparison pooled across the three genes *CALD1*, *KATNA1, SLITRK1*. Genes and treatments were treated as random and fixed effect, respectively. P values are adjusted for multiple comparison (** P ≤ 0.01, *** P ≤ 0.001). Overall, there was a clear treatment effect (F(12, 24)=32.954, P ≤ 0.001).

To test if the CRISPY mix increases TNS in other human pluripotent stem cell lines we edited the gene *KATNA1* in SC102A1 iPSCs and H9 ESCs using Cas9n plasmid nucleofection. TNS increased 2.6-fold and 2.8-fold, respectively, and the increase was bigger than when using NU7026 alone (Fig. 3C). When we used Cas9, which introduces blunt-ended DSBs, no significant effect was seen when adding other small molecules in addition to NU7026 (Fig. 3B). In contrast, the CRISPY mix together with Cpf1 ribonucleoprotein, which produces staggered DNA cuts, introduced by electroporation in 409-B2 hiPSCs increased TNS 2.9-fold for *HPRT* and 4-fold for *DMNT1* (Fig. 3D). Addition of only NU7026 increased TNS 2.1-fold for *HPRT* and 2.4-fold for *DMNT1*.

Thus, NU7026, Trichostatin A, MLN4924 and NSC 15520 increase TNS with Cas9n and Cpf1 when applied either singly or together (Fig. 2A and 3A, C and D). NU7026 inhibits DNA-PK (Fig. 1), a major complex in NHEJ pathway^1^ and has been shown to increase knock-in efficiency in hiPSCs^22^. Trichostatin A activates an ATM-dependent DNA damage signaling pathway^26^. MLN4924 inhibits the Nedd8 activating enzyme (NAE) and has been shown to inhibit the neddylation of CtIP which leads to an increase of the extent of DNA end resection at strand breaks thereby promoting HDR^27^ by leaving ssDNA stabilized by RPA that can undergo recombination. NSC15520 prevents the association of RPA with p53 and RAD9^28, 29^, possibly increasing the abundance of RPA available which could favor HDR. RS-1, SCR7 and L755507 for which there are conflicting reports on their capacity to increase precise genome editing (Supplementary Table 1) showed no measurable effect in our hands on TNS neither in the Cas9 or the Cas9n cells.

It has been suggested that different types of cuts engage different repair pathways, because 5’ overhanging ends yield higher levels of HDR than 3’ overhangs^5^. The fact that Trichostatin A and MLN4924 increase TNS with Cas9n and Cpf1 but has no effect with Cas9 suggests that blunt ends introduced by Cas9 may be repaired by different mechanisms versus 5’ overhanging DNA ends introduced by Cpf1 or Cas9n. While RAD51 is obviously important for classical homologous recombination with dsDNA^1^ it is possible that RAD52, rather than RAD51, could be responsible for HDR with ssDNA donors since RAD52 is needed for annealing of ssDNA^4^. Our results of the effect of RAD52 inhibitor AICAR, RAD51 inhibitor B02, and RAD51 enhancer RS-1 on TNS efficiency suggests that RAD51 and not RAD52 is important for precise editing with ssODN, since inhibiton of RAD51 by B02 halved and inhibiton of RAD52 had no effect on TNS efficiency. Interestingly, RS-1 had no beneficial effect on TNS.

In summary, we show that in an inducible Cas9n iPSC line and using the optimized protocol described we achieve TNS and indels in more than 90% of chromosomes (Fig. 3A). Using the CRISPY mix of small molecules we achieved almost 50% TNS in hiPSCs (Fig. 3A), to our knowledge the highest efficiency of precise genome editing in human pluripotent stem cells described to date.

## Acknowledgements

We would like to thank Malgorzata Gac for assistance with FACS-sorting, Anna Kirstein for experimental help in validation of the iCRISPR cell lines and Roger Mundry for help with statistical analysis. Furthermore, we would like to thank Antje Weimann and Barbara Höber for DNA sequencing and Svante Pääbo for comments on the manuscript and helpful discussions. This work was supported by the Max Planck Society.

## Author contributions

S.R. conceived the idea, S.R. and T.M planned the experiments and S.R. performed the experiments. S.R. and T.M wrote the paper.

## Online Methods

### Cell culture

iPSC lines cultured for this project included human 409-B2 iPSC (female, Riken BioResource Center) and SC102A1 iPSC (male, BioCat GmbH), as well as H9 ESC (female, WiCell Research Institute, Ethics permit AZ 3.04.02/0118). Cells were grown on Matrigel Matrix (Corning, 35248) at 37°C in a humidified incubator gassed with 5% CO_2_. mTeSR1 (StemCell Technologies, 05851) with mTeSR1 supplement (StemCell Technologies, 05852) was used as culture media. Media was replaced every day. Cell cultures were maintained 4-6 days until ∼ 80% confluency, dissociated using EDTA (VWR, 437012C) and subcultured at a 1:6 to 1:10 dilution. The media was supplemented with 10 μM Rho-associated protein kinase (ROCK) inhibitor Y-27632 (Calbiochem, 688000) after cell splitting for one day in order to increase cell survival.

### Generation and validation of iCRISPR cell lines

Human 409-B2 iPSCs were used to create an iCRISPR-Cas9 line as described by Gonzalez et al. ^11^ (GMO permit AZ 54-8452/26). For the production of iCRISPR-Cas9n line Puro-Cas9 donor was subjected to site-directed mutagenesis with the Q5 mutagenesis kit to introduce the D10A mutation (New England Biolabs, E0554S). Primers were ordered from IDT (Coralville, USA) and are shown in Supplementary Table 3. Expression of the pluripotency markers SOX2, OCT-4, TRA1-60 and SSEA4 in iCRISPR lines was validated using the PSC 4-Marker immunocytochemistry kit (Molecular Probes, A24881) (data not shown). Quantitative PCR was used to confirm doxycycline inducible Cas9 or Cas9n expression and digital PCR was used to exclude off-target integration of the iCRISPR cassettes (data not shown).

### Small molecules

Commercially available small molecules used in this study were NU7026 (SIGMA, T8552), Trichostatin A (SIGMA, T8552), MLN4924 (Adooq BioScience, A11260), NSC 19630 (Calbiochem, 681647), NSC 15520 (ChemBridge, 6048069), AICAR (SIGMA, A9978), RS-1 (Calbiochem, 553510), Resveratrol (Selleckchem, S1396), SCR7 (XcessBio, M60082-2s), L755507 (TOCRIS, 2197), B02 (SIGMA, SML0364) and STL127685 (Vitas-M). STL127685 is a 4-fluorophenyl analog of the non-commercially available STL127705. Stocks of 15mM (or 10mM for NU7026) were made using dimethylsulfoxide (DMSO)(Thermo Scientific, D12345). Solubility is a limiting factor for NU7026 concentration. Suitable working solutions for different concentrations were made so that addition of each small molecule accounts for a final concentration of 0.08% (or 0.2% for NU7026) DMSO in the media. Addition of all small molecules would lead to a final concentration of 0.7% DMSO.

### Design of gRNAs and ssODNs

We chose to introduce one desired mutation in three genes *CALD1*, *KATNA1* and *SLITRK1* back to the state of the last common ancestor of human and Neanderthal^30^. gRNA pairs for editing with the Cas9n nickase were selected to cut efficiently at a short distance from the desired mutation and from the respective partnering sgRNA. The efficiency was estimated with the sgRNA scorer 1.0 tool^31^ as a percentile rank score. gRNAs for nickase editing were designed to have the desired mutation and Cas9-blocking mutations to prevent re-cutting of the locus and had 50nt homology arms upstream and downstream of each nick (Supplementary Fig. 2). gRNA of the nickase gRNA pair that cuts closer to the desired mutation was used for Cas9 nuclease editing together with a 90nt ssODN centered at the desired mutation and containing a Cas9-blocking mutation (Supplementary Fig. 2). ssODNs for editing of *HPRT* and *DMNT1* using Cpf1 were designed to contain a blocking mutation near the PAM site and an additional mutation near the cut. gRNAs (crRNA and tracR) and ssODN were ordered from IDT (Coralville, USA). ssODNs and crRNA targets are shown in Supplementary Table 3.

### Lipofection of oligonucleotides

Cells were incubated with media containing 2μg/ml doxycycline (Clontech, 631311) 2 days prior to lipofection. Lipofection (reverse transfection) was done using the alt-CRISPR manufacturer’s protocol (IDT) with a final concentration of 7.5nM of each gRNA and 10nM of the respective ssODN donor. In brief, 0.75μl RNAiMAX (Invitrogen, 13778075) and the respective oligonucleotides were separately diluted in 25μl OPTI-MEM (Gibco, 1985-062) each and incubated at room temperature for 5min. Both dilutions were mixed to yield 50μl of OPTIMEM including RNAiMAX, gRNAs and ssODNs. The lipofection mix was incubated for 20-30min at room temperature. During incubation cells were dissociated using EDTA for 5min and counted using the Countess Automated Cell Counter (Invitrogen). The lipofection mix, 100μl containing 25.000 dissociated cells in mTeSR1 supplemented with Y-27632, 2μg/ml doxycycline and the respective small molecule(s) to be tested were througoutly mixed and put in one well of a 96 well covered with Matrigel Matrix (Corning, 35248). Media was exchanged to regular mTeSR1 media after 24hours.

### Ribonucleoprotein nucleofection

The recombinant *A.s.* Cpf1 protein and electroporation enhancer was ordered from IDT (Coralville, USA) and nucleofection was done using the manufacturer’s protocol, except for the following alterations. Nucleofection was done using the B-16 program of the Nucleofector 2b Device (Lonza) in cuvettes for 100 μl Human Stem Stell nucleofection buffer (Lonza, VVPH-5022) containing 1 million 409-B2 iPSCs, 78pmol electroporation enhancer, 0.3nmol gRNA, 200pmol ssODN donor, 252pmol Cpf1. Cells were counted using the Countess Automated Cell Counter (Invitrogen).

### FACS-sorting

Introduction of 2μg plasmid DNA (pSpCas9n(BB)-2A-GFP (PX461), Addgene #48140) into cells not expressing Cas9 inducably was done using the B-16 program of the Nucleofector 2b Device (Lonza) in cuvettes for 100 μl Human Stem Stell nucleofection buffer (Lonza, VVPH-5022) containing 1 million of either SC102A1 iPSC or H9 ESC. Cells were counted using the Countess Automated Cell Counter (Invitrogen). 24h after nucleofection cells were dissociated using Accutase (SIGMA, A6964), filtered to obtain a single cell solution, and subjected to fluorescence associated cell sorting (FACS) for GFP expressing cells. During sorting with the BD FACSAria III (Becton-Dickinson) cells were kept at 4°C in mTeSR1 supplemented with Y-27632. 48h after sorting cells were subjected to lipofection with sgRNAs, ssODNs and treatment with small molecules.

### Illumina library preparation and sequencing

Three days after lipofection cells were dissociated using Accutase (SIGMA, A6964), pelleted, and resuspended in 15μl QuickExtract (Epicentre, QE0905T). Incubation at 65°C for 10min, 68°C for 5min and finally 98°C for 5min was performed to yield ssDNA as a PCR template. Primers for each targeted loci containing adapters for Illumina sequencing were ordered from IDT (Coralville, USA) (see Supplementary Table 3). PCR was done in a T100 Thermal Cycler (Bio-Rad) using the KAPA2G Robust PCR Kit (Peqlab, 07-KK5532-03) with supplied buffer B and 3 μl of cell extract in a total volume of 25μl. The thermal cycling profile of the PCR was: 95°C 3min; 34x (95° 15sec, 65°C 15sec, 72°C 15sec); 72°C 60sec. P5 and P7 Illumina adapters with sample specific indices were added in a second PCR reaction^32^ using Phusion HF MasterMix (Thermo Scientific, F-531L) and 0.3μl of the first PCR product. The thermal cycling profile of the PCR was: 98°C 30sec; 25x (98° 10sec, 58°C 10sec, 72°C 20sec); 72°C 5min. Amplifications were verified by size separating agarose gel electrophoresis using EX gels (Invitrogen, G4010-11). The indexed amplicons were purified using Solid Phase Reversible Immobilization (SPRI) beads^33^. Double-indexed libraries were sequenced on a MiSeq (Illumina) giving paired-end sequences of 2 x 150 bp. After base calling using Bustard (Illumina) adapters were trimmed using leeHom^34^.

### CRISPResso analysis

CRISPresso^35^ was used to analyse sequencing data from CRISPR genome editing experiments for percentage of wildtype, targeted nucleotide substitutions (TNS), indels and mix of TNS and indels. Parameters used for analysis were ‘-w 20’, ‘--min_identity_score 70’ and ‘--ignore_substitutions’ (analysis was restricted to amplicons with a minimum of 70% similarity to the wildtype sequence and to a window of 20bp from each gRNA; substitutions were ignored, as sequencing errors would be falsly characterized as NHEJ-events)

### Statistical analysis

Significances of changes in TNS efficiencies were determined using a two-way-ANOVA and Tukey multiple comparison pooled across the three genes *CALD1*, *KATNA1*, *SLITRK1*. Genes and treatments were treated as random and fixed effect, respectively. Hence, we tested the effect of treatment against its interaction with gene^36^. Analysis included 3 technical replicates for each gene. We checked for whether the assumptions of normally distributed and homogeneous residuals were fulfilled by visual inspection of a QQ-plot^37^ and residuals plotted against fitted values^38^. These indicated residuals to be roughly symmetrically distributed but with elongated tails (i.e., too large positive and negative residuals) and no obvious deviations from the homogeneity assumption. P values are adjusted for multiple comparison. Statistical analysis was done using R.

